# A Higher Proportion of Craniosynostosis Genes Are Cancer Driver Genes

**DOI:** 10.1101/872093

**Authors:** Suchir Misra, Andrew Shih, Xiao-Jie Yan, Wentian Li

**Affiliations:** The Robert S. Boas Center for Genomics and Human Genetics, The Feinstein Institutes for Medical Research, Northwell Health, Manhasset, NY, USA; The Karches Center for Chronic Lymphocytic Leukemia Research, The Feinstein Institutes for Medical Research, Northwell Health, Manhasset, NY, USA

**Keywords:** craniosynostosis, cancer driver genes, gene-set, enrichment analysis, signaling pathways, oncogenesis recapitulates ontogenesis

## Abstract

Craniosynostosis (CRS) is a congenital abnormality deformity with a heterogenous genetic contribution. Previously, there are two attempts to collect genes that are genetically associated with craniosynostosis and some related syndromes with 57 (Twigg and Wilkie, 2015) and 39 (Goos and Mathijssen, 2019) genes identified, respectively. We expanded this list of craniosynostosis genes by adding another 17 genes with an updated literature search. These genes are shown to be more likely to be intolerant to functional mutations. Of these 113 craniosynostosis genes, 21 (19% vs. 1.5% baseline frequency) are cancer driver genes, a 14-fold enrichment. The cancer-craniosynostosis connection is further validated by an over-representation analysis of craniosynostosis genes in KEGG cancer pathway and several cancer related gene-sets. Many cancer-craniosynostosis overlapping genes participate in intracellular signaling pathways, which play a role in both development and cancer. This connection can be viewed from the oncogenesis recapitulates ontogenesis framework. Nineteen craniosynostosis genes are transcription factor genes (16.8% vs. 8.2% baseline), and craniosynostosis genes are also enriched in targets of certain transcription factors or micro RNAs.

## Introduction

Craniosynostosis is a congenital condition when the suture of a baby’s skull does not close in a timely fashion, causing abnormal skull shape (Kabbani and Raghuveer, 2004). This premature fusion of sutures may also lead to other abnormal traits, a situation called a craniosynostosis syndrome (Muenke and Wilkie, 2014; Ko, 2016). There are 100-200 syndromes associated with craniosynostosis, including Apert, Pfeiffer, Crouzon, Antley-Bixler, Saethre-Chotzen, and Muenke syndromes (Wilkie et al., 1995; Carinci et al., 2005); at least 15 of the syndromes are listed in Online Mendelian Inheritance in Man (*www.omim.org*), and close to 40 are listed in National Organization for Rare Disorders (*rarediseases.org*). Craniosynostosis is highly heterogeneous in its potential causes, etiology of individual cases is often unknown (Wilkie, 1997), and has long been known to be a multifactorial disease (Hunter and Rudd, 1976). Through genetic studies, tens of genes are shown to be associated with craniosynostosis (Twigg and Wilie, 2015), providing a foundation for further elucidation of the mechanism of the trait. Like any phenotypes, there should also be environmental contributions to craniosynostosis (Shashi and Hart, 2002; Zeiger et al., 2002; Durham et al., 2017), illustrated by an observation of identical twins with discordant craniosynostosis (Magge et al., 2017).

If a specific gene is part of a known biological pathway, the genes function and its role in a disease might be hypothesized. However, due to the extreme heterogeneity of craniosynostosis, no single pathway stands out as the major pathway for all cases (Twigg and Wilie, 2015). We adopt a different strategy to treat the collection of genes that are genetically associated with craniosynostosis as a single unit (a gene-set, a molecular signature, etc.) (Jürgens, 1985; Subramanian et al., 2005; Li et al., 2015b). One gene-set can be checked with any other gene-sets for overlapping of common genes. If the two gene-sets share more common genes than expected by chance, an over-representation or enrichment is claimed, which provides evidence that the two gene-sets are biologically related (Khatri et al., 2012).

Naturally, the major focus on the molecular understanding of craniosynostosis is on early embryonic developmental pathways, such as the fibroblast growth factor (FGF) signaling pathway (Ornitz and Marie, 2002) that is associated with several downstream transduction pathways of RAS/MAPK (Takenouchi et al., 2014), PI3K/AKT (Dufour et al., 2008), PLC*γ* (Moenning et al., 2008), STAT (Lokau and Garbers, 2019), etc. The link between craniosynostosis and cancer was first suggested by the observation that a craniosynostosis-associated gene, FGFR2, experiences a higher than expected somatic mutation rate in endometrial cancer (Pollock et al., 2007).

In this paper, we aim at extending the craniosynostosis-cancer connection from a single gene (i.e., FGFR2) to the level of pathways and gene-sets. Towards this goal, we would like to have a more up-to-date list of genes that are genetically associated with craniosynostosis (i.e., genes that contain, uniquely or preferably, germline mutations in craniosynostosis patients). Previously, there have been two attempts to collect craniosynostosis genes: one in 2015 with 57 genes (Twigg and Wilie, 2015) and a more recent one in 2019 with 39 genes (Goos and Mathijssen, 2019).

Most of these genes are identified through a combination of family studies, candidate gene investigation, mouse model validation, etc. Recently, there are new strategies, such as GWAS (genome-wide association study) on common genetic variants (Justice et al., 2012), and exome sequencing followed by summing all rare variants (Timberlake et al., 2017). Without distinguishing different gene mapping methods, we proceed by a systematic search of all PUBMED titles/abstracts that contain the word of “craniosynostosis and “genetic, finding all gene names, then followed by a manual check of the original literature. An extra 17 genes have been added to the craniosynostosis genes list.

On the other hand, a list of cancer driver genes (i.e., somatic mutations in these genes are positive selected in cancer cell clonal expansion) is readily available with 125 genes in a 2013 study (Vogelstein et al., 2013), and 299 genes in a more recent analysis (Tokheim et al., 2016). We compare our craniosynostosis gene list and cancer driver gene lists to examine if the overlap between them is more than expected by chance.

The result section below is organized by a description of our compilation of craniosynostosis genes; the observation that these genes are more intolerant to functional mutations; these genes are enriched in cancer driver gene lists; an enrichment analysis of these genes in all gene-set from a gene-set database; an enrichment analysis of these genes in the Ingenuity pathway database; and their enrichment in the transcription factor list. The result section is followed by the Method/Data section and Discussion/Conclusion section.

## Results

### Compilation of a more complete list of craniosynostosis genes

In 2015, a set of 57 genes are summarized as being genetically associated with craniosynostosis (CRS) (Twigg and Wilie, 2015), i.e., variants/mutations are observed in these genes in CRS patients that are less likely to be observed in people without CRS. A more recent paper adds 39 more CRS genes published from 2015-2017 (Goos and Mathijssen, 2019), resulting in a total of 96 genes. The reason for so many genes being linked to the disease can be understood by the perspective of heterogeneity (McClellan and King, 2010).

When some public PUBMED-search programs such as PolySearch (*polysearch.ca*) (Liu et al., 2015) or HuGE Literature Finder (*phgkb.cdc.gov/PHGKB/startPagePubLit.actio*) (Lin et al., 2006) were used, we failed to find any new CRS genes. The reason is that these programs are designed to reduce the number of false positives, aiming at a shorter list of genes, not longer. As these public domain programs did not serve our purposes, we designed our own pipeline via a PUBMED search.

In our approach, we first filtered all PUBMED titles/abstracts by the MeSH term of craniosynostosis and genetic. Then, a list of HUGO approved human gene names was used to find any matching abbreviations in the abstract (two-letter items are ignored). With this raw list of potential gene names, we manually checked each item and the corresponding abstract to determine the appropriateness of the match.

These 17 new genes found through this process were added to the CRS genes list: AXIN2 (Yilmaz et al., 2018), BBS9 (Justice et al., 2012; Sewda et al., 2019), BCOR (O’Byrne et al., 2017) (for Oculo-facio-cardio-dental syndrome, or microphthalmia syndrome), BGLAP (Sowińska-Seidler et al., 2018), COLEC10 (for 3MC syndrome) (Munye et al., 2017), FGFRL1 (Rieckmann et al., 2009) (for Antley-Bixler syndrome), GCK (for Greig cephalopolysyndactyly syndrome) (Zung et al., 2011), LMNA (Sowińska-Seidler et al., 2018), PPP3CA (Mizuguchi et al., 2018), PTH2R (Kim et al., 2015), RAF1 (for Noonan syndrome with multiple lentigines, or leopard syndrome) (Rodríguez et al., 2019), SIX2 (for frontonasal dysplasia syndrome) (Hufnagel et al., 2016), SMURF1, SPRY1, SPRY4 (Timberlake et al., 2016, 2017), TCOF1 (for Treacher Collins syndrome) (Horiuchi et al., 2004), TNFRSF11B (for Juvenile Paget disease) (Saki et al., 2013).

There can be various reasons why the above genes are not included in the two collections by Twigg-Wilkie and Goos-Mathijssen. Obviously, if the paper was published after 2018, it would not be in the previous lists. Also, we included gene mutations in rare syndromic cases which include CRS as one of many phenotypes. Note that we did not consider genes causing CRS in animal models (e.g., BMP3 (Schoenebeck et al., 2012), DUSP6 (Li et al., 2007), SNAI1, SNAI2 (Oram and Gridley, 2005)). We also did not consider genes that are differentially expressed in relevant tissues in CRS (e.g., FGF7, SFRP4, VCAM1 (Zhang et al., 2002; Stamper et al., 2011), NELL1 (Zhang et al., 2002), IGF1 (Al-Rekabi et al., 2016)). These genes may still play a role in the etiology of CRS, or, they might be discovered to harbor germline mutations in the future, but are excluded for now to be consistent with the previous lists.

The three compilations ((Twigg and Wilie, 2015), (Goos and Mathijssen, 2019), and here) add up to 113 genes that are genetically associated with craniosynostosis. These 113 craniosynostosis genes are: ABCC9, ADAMTSL4, AHDC1, ALPL, ALX4, ASXL1, ATR, AXIN2, B3GAT3, BBS9, BCOR, BGLAP, BMP2, BRAF, CD96, CDC45, CHST3, COLEC10, COLEC11, CRTAP, CTSK, CYP26B1, DPH1, EFNA4, EFNB1, ERF, ESCO2, FAM20C, FBN1, FGF9, FGFR1, FGFR2, FGFR3, FGFRL1, FLNA, FREM1, FTO, GCK, GLI3, GLIS3, GNAS, GNPTAB, GPC3, HNRNPK, HUWE1, IDS, IDUA, IFT122, IFT140, IFT43, IGF1R, IHH, IL11RA, IL6ST, IRX5, JAG1, KANSL1, KAT6A, KAT6B, KMT2D, KRAS, LMNA, LMX1B, LRP5, MASP1, MED13L, MEGF8, MSX2, NFIA, NTRK2, OSTM1, P4HB, PHEX, POR, PPP1CB, PPP3CA, PTH2R, PTPN11, PTPRD, RAB23, RAF1, RECQL4, RSPRY1, RUNX2, SCARF2, SCN4A, SEC24D, SH3PXD2B, SHOC2, SIX2, SKI, SLC25A24, SMAD6, SMC1A, SMO, SMURF1, SOX6, SPECC1L, SPRY1, SPRY4, STAT3, TCF12, TCOF1, TGFBR1, TGFBR2, TMCO1, TNFRSF11B, TWIST1, WDR19, WDR35, ZEB2, ZIC1, and ZNF462.

More detailed information on the newly added 17 CRS genes are summarized in the Appendix. The gene OSTM1 is labeled as OSTEM1 in (Mahmoud Adel et al., 2013; Goos and Mathijssen, 2019) which is not a HGNC approved name. Also note that BGLAP and LMNA in our new addition are within a 1.26 Mb deletion and each gene was hypothesized as causing the disease (Sowińska-Seidler et al., 2018).

### Craniosynostosis genes tend to be intolerant to functional mutations

Sequencing a large number of normal samples allows us to learn which genes tend to have more amino-acid-changing and/or loss-of-function mutations and which genes tend to have less. The former is called “tolerant” and the latter “intolerant”. We tested three measures of intolerance with two goals in mind. One is to check whether our CRS gene list as a whole tend to be intolerant to functional mutations. Another is to find out which genes in the list are more tolerant to mutations than others, which flag them as potential false positives.

The RVIS (Residual Variation Intolerance Score) (Petrovski et al., 2013) is derived from the regression of sum of all common (major allele frequency larger than 0.001) functional variants within a gene (Y) over sum of all positions with variant in the gene (X). The residual (difference between the observed Y and the expected value from the regression line), standardized by the standard deviation, is RVIS (Petrovski et al., 2013). Genes with negative (positive) are intolerant (tolerant) to functional mutations. Of 104 CRS genes (out of 113) with RVIS values, 82 are negative (and 19 are positive). The percentage of intolerant CRS genes is 79%, as compared to the baseline percentage of 53% (Fisher’s test p-value= 9 × 10^−9^). Direct comparison of RVIS in CRS gene list and outside also leads to a significant difference: the t-test p-value is 3.7 × 10^−9^ and the Mann-Whitney-Wilcoxon test p-value is 1.6 × 10^−11^.

The LOEUF (Loss-of-function Observed-over-Expected Upper bound Fraction) is based on a coarse-grained version of the regression of observed loss-of-function variants (Y) over expected (X) (Karczewski et al., 2019), for data from the normal individuals. A large (small) LOEUF value implies a tolerant (intolerant) gene. Of the 113 CRS genes, 95 have LOEUF smaller than 1 (84.1%). This percentage is significantly larger than the baseline proportion (55.5%) (Fisher’s test p-value = 1.3 × 10^−10^). Direct comparison of LOEUF between CRS genes and the rest leads to the same conclusion (Mann-Whitney-Wilcoxon test p-value is 2.4 × 10^−16^; t-test is not used because the distribution of LOEUF is not normal).

The GDI (Gene Damage Index) is calculated by summing all variants’ predicted damage impact score according to CADD (Combined Annotation Dependent Depletion) (Kircher et al., 2014) in a normal dataset (Itan et al., 2015). A large (small) GDI implies a tolerant (intolerant) gene. A comparison of the GDI score itself between CRS genes and other genes does not lead to any significant results. In a recent assessment of various variant pathogenicity prediction programs, CADD is not performing well (Niroula and Vihinen, 2019). It may cast doubt on the usefulness of raw GDI score. However, if we use the categorical label on whether a gene has LOW, MEDIUM, or HIGH damage provided by the GDI site, genes showing high level of damaging mutations in normal samples might be potential false positives. According to the assessment from this approach, these genes in the CRS list: FREM1, KANSL1, ALX4, RECQL4, may need further examination. Despite the possibility that some genes in the list may be false positives, as a whole, we show evidence that these genes are intolerant to functional mutations.

### Enrichment of craniosynostosis genes in the list of cancer driver genes

A cancer driver gene is defined as those whose mutations “increase net cell growth under the specific microenvironmental conditions that exist in the cell in vivo” (Tokheim et al., 2016), or genes whose mutations “promote tumorigenesis” (Vogelstein et al., 2013). Many cancer driver genes prediction relies on the non-random mutation patterns, instead of mutation frequencies. For example, the 20/20 rule states that for oncogenes, > 20% of somatic mutations should occur in recurrent positions and to be missense, whereas for tumor suppressing genes, 20% of mutations should be inactivating (loss-of-function, truncating) (Vogelstein et al., 2013).

We obtained the list of 299 cancer driver genes from (Bailey et al., 2018), which is decided by a combination of 26 computer programs. We first tested enrichment of cancer driver genes individually in Twigg-Willie’s list, in Goos-Mathijssen’s list, and in our newly added list. The results are shown in Table 1. There are 12 (out of 57), 6 (out of 39), 3 (out of 17) CRS genes that are in the cancer driver gene list, for the three collections. The proportions are 21%, 15%, and 18% respectively. On the other hand, the baseline proportion of cancer driver genes can be simply 299/20000 or 1.5%. Clearly, cancer driver genes are greatly enriched in the three CRS gene lists. The overrepresentation test (using Fisher’s test) leads to p-values of 6 × 10^−11^, 3 × 10^−5^ and 2 × 10^−3^, respectively.

**Table 1:** Enrichment of cancer driver genes (n=299 set from (Bailey et al., 2018) and n=125 set from (Vogelstein et al., 2013)) in various lists of craniosynostosis genes ((Twigg and Wilie, 2015; Goos and Mathijssen, 2019) and this paper). The 299 cancer driver genes are used to compare with the craniosynostosis genes. for cancer driver genes frequency is 299/20000= 1.5% or 125/20000= 0.6%, respectively in the two cancer driver gene lists, assuming the total number of human genes to be 20000.

|  | n=299 driver |  |  |  | n=125 driver |  |  |
| --- | --- | --- | --- | --- | --- | --- | --- |
| CRS gene-set | N | $N_{dri}$ (per) | odds-ratio | Fisher p-value | $N_{dri}$ (per) | OR | pv |
| Twigg,Wilkie(-2015) | 57 | 12(21%) | 17.6 | $6 \times 10^{-11}$ | 5 (8.8%) | 15.3 | $3 \times 10^{-5}$ |
| core genes | 20 | 5 (25%) | 22.0 | $1 \times 10^{-5}$ | | | |
| Goos,Mathijssen(2015-2017) | 39 | 6 (15%) | 12.0 | $3 \times 10^{-5}$ | 3 (7.7%) | 13.3 | $2 \times 10^{-3}$ |
| multiple-per | 22 | 2 (9%) | 6.6 | 0.04 |  |  |  |
| single-per | 17 | 4 (24%) | 20.3 | $1 \times 10^{-4}$ | | | |
| Twigg+Goos | 96 | 18 (19%) | 15.2 | $1 \times 10^{-14}$ | 8 (8.3%) | 14.5 | $2 \times 10^{-7}$ |
| Suchir et al (-2019) | 17 | 3 (18%) | 14.1 | 0.002 | 1 (5.9%) | 9.9 | 0.1 |
| All: Twigg + Goos + Suchir | 113 | 21 (19%) | 14.7 | $1.2 \times 10^{-16}$ | 9 (8%) | 13.8 | $7 \times 10^{-8}$ |

Within Twigg-Wilkie’s list, there are 20 CRS genes are labeled as core genes. And 5 of these 20 genes are cancer driver genes (25%). The Goos-Mathijssen’s list can be split into two sublists: one with evidence of mutation in multiple persons, and another with mutation observed only on one patient. The cancer driver genes are in 9% and 24% of the two sub-lists. All these proportions are much higher than the 1.5% expected by chance. If we combine the three CRS gene collections (total 113 genes), 21 (or 19%) are cancer driver genes, with the p-value of 1.2 × 10^−16^.

To address the question on whether the significant enrichment of cancer driver genes in CRS genes is robust, with respect to the compilation of driver genes itself, we re-ran the analysis using an early cancer driver gene list (125 genes) (Vogelstein et al., 2013). The result is summarized in Table 1 (right). The baseline proportion of this cancer driver genes is 125/20000 = 0.6%, whereas the proportion of them in the three CRS gene lists are 8.8%, 7.7%, and 5.9%. The corresponding Fisher’s test p-values are 3 × 10^−5^, 2 × 10^−3^, and 0.1. When all three CRS gene collections are combined, 9 out of 113 (8%) are cancer driver genes, a significant enrichment (p-value= 7×10^−8^). These results firmly establish the fact that cancer driver genes are over-presented in CRS genes.

Our results can be made more precise by using a more accurate number of human genes (less than 20000), and/or by excluding CRS genes from the comparison group. These changes do not affect the conclusion (result not shown). The 21 cancer driver genes that match the CRS gene list are ASXL1, ATR, AXIN2, BCOR, BRAF, FGFR1, FGFR2, FGFR3, FLNA, GNAS, HUWE1, IL6ST, KANSL1, KMT2D, KRAS, PTPN11, PTPRD, RAF1, SMC1A, TCF12, and TGFBR2.

### Enrichment analysis of craniosynostosis genes for all gene-sets in MSigDB

In order to further examine the craniosynostosis-cancer connection, we compared the 113-CRS-gene list with all gene-sets in MSigDB (Liberzon et al., 2015). We checked whether some genes in one list may never have a chance to be present in the second list, due to the two-gene-universe situation (e.g. (Swaminathan and Fury, 2012)). Indeed, MSigDB v6.2 has not updated all gene names to the current standard, with KANSL1 written as KIAA1267, and KMT2D written as MLL2 in MSigDB. We manually corrected these inconsistent gene names. Similarly, non-coding genes in MSigDB may never have a chance to match our CRS gene list which is all protein-coding. Therefore, only protein-coding genes are kept in MSigDB gene-sets in this enrichment analysis. The C1 MSigDB collection is chromosome location based gene clusters, which is of no interest to us and thus not used here.

The Hallmark, C4, C6, and C7 collection do not have any gene-sets to be significantly overlapping with CRS gene list at the family-wide significance p-value of = 5.7 × 10^−7^. However, when using the collections-specific multiple testing correction, the TGF-beta signaling (Hallmark) (*p* = 3.3 × 10^−5^, OR=15.4) is significant. This pathway has already been discussed as being linked to CRS (Opperman et al., 1997; Hunenko et al., 2001) (see Table 2 for matching genes, and Fig.1(D) for the volcano plot (Li, 2012)).

**Figure 1:**
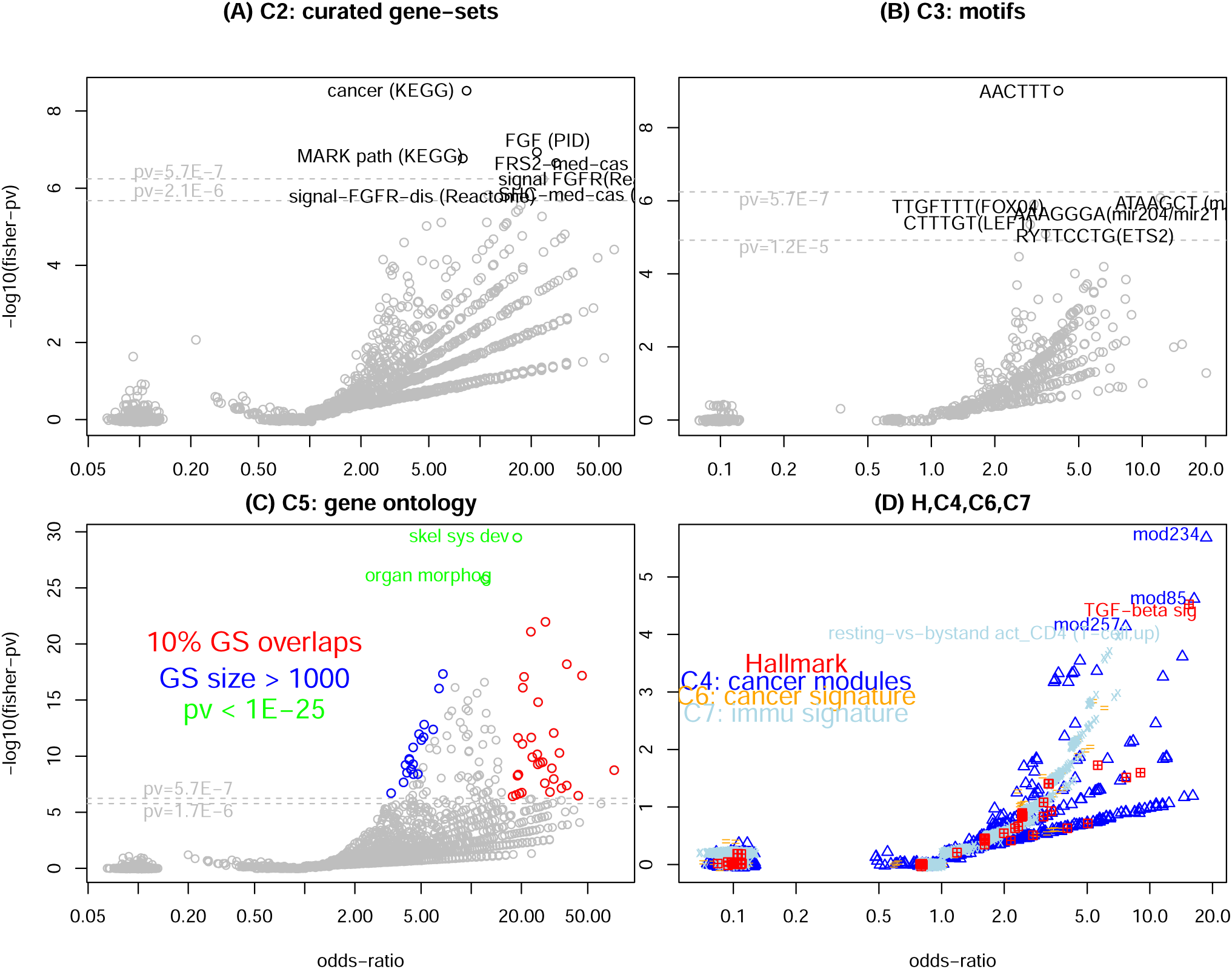
Volcano plot (x: (log) odds-ratio, y: −log_10_(Fisher p-value) (Li et al., 2014)) for enrichment (over-representation) analysis of craniosynostosis genes in MSigDB gene-sets: (A) curated gene-sets, including KEGG, Reactome, PID (C2); (B) motif based gene-sets (C3); (C) gene ontology gene-sets (C5). Two groups of significantly enriched gene-sets are highlighted: large gene-sets (> 1000 genes) (in blue), and high proportion (> 10%) of overlapping genes (in red); (D) other categories of gene-sets (hallmark, microarray expression derived signatures (C4, C6, C7)).

**Table 2:**
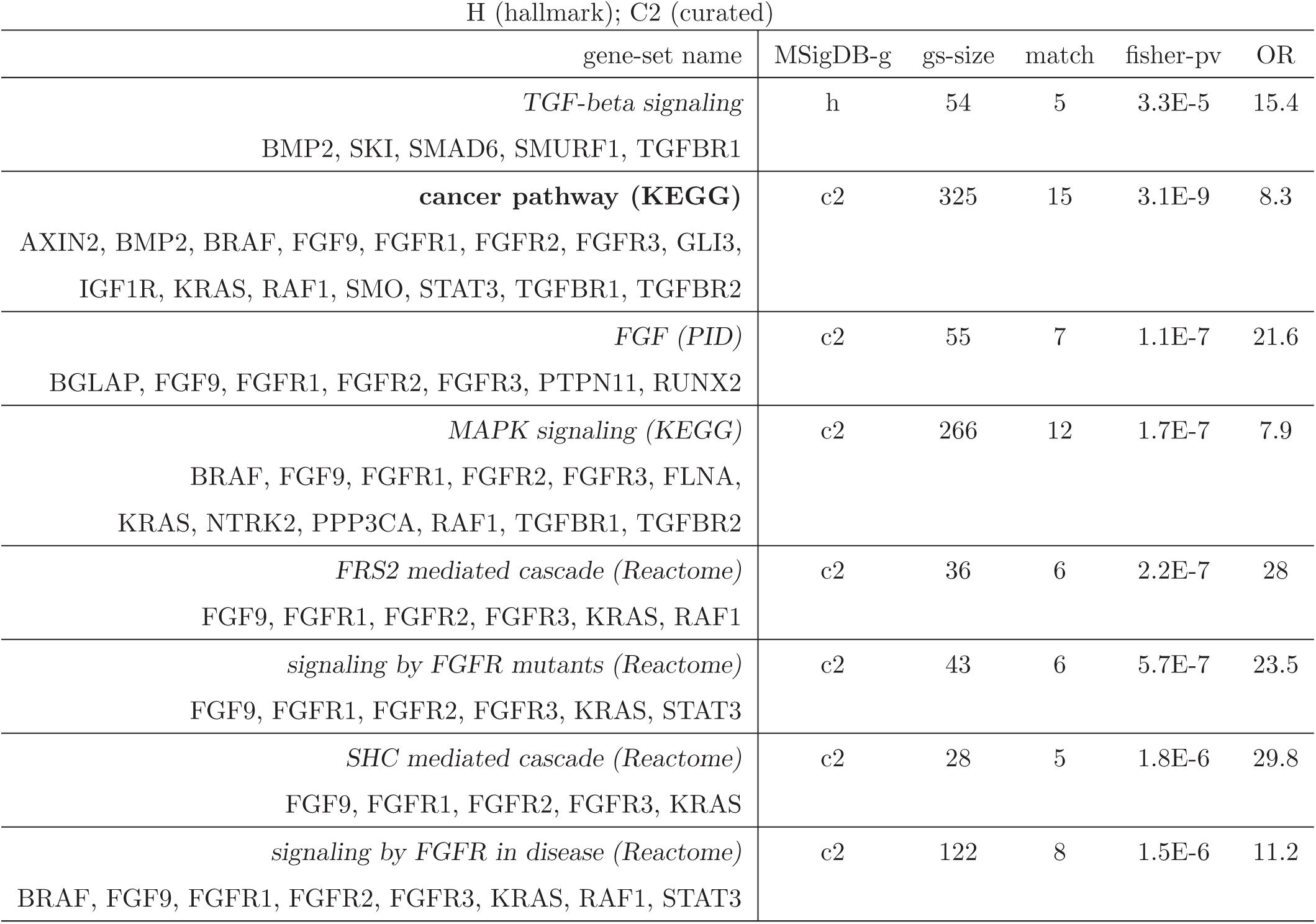
Gene-sets in MSigDB H and C2 that are enriched with craniosynostosis genes (significant at the family-wide level), and the corresponding overlapping gene names. Gene-set size, overlapping set size, Fisher’s test p-value and odds-ratio (OR) are also listed.

The C2 gene-set collection in MSigDB includes some other popular pathways and gene-sets databases such as KEGG (Kanehisa and Sato, 2000) and REACTOME (Fabregat et al., 2018). Using a stringent criterion of (Bonferroni correction) p-value < 0.01*/n*_*GS*_, which is 5.7 × 10^−7^ if we use *n*_*GS*_ = 17484 for all gene-sets in MSigDB (used in the enrichment analysis), or 2.1 × 10^−6^ if we only count the number of gene-sets in C2 (*n*_*GS*_ = 4762), 3 or 5 gene-sets are family-wise significant. These top five gene-sets are: cancer pathway (KEGG) (p-value (*p*) = 3.1 × 10^−9^, odds-ratio (OR) =8.3), FGF pathway (defunct PID (Schaefer et al., 2009)) (*p* = 1.1 × 10^−7^, OR=21.6), MAPK signaling pathway (KEGG) (*p* = 1.7 × 10^−7^, OR=7.9), FRS2 mediated cascade (REACTOME) (*p* = 2.2 × 10^−7^, OR=28), and signaling by FGFR mutants (REACTOME) (*p* = 5.7 × 10^−7^, OR=29.8), These results and matching genes are listed in Table 2, and volcano plot (Li, 2012) in Fig.1(A).

The involvement of FGF (fibroblast growth factor), FGFR (fibroblast growth factor receptor), and MAPK/ERK signaling pathways in CRS has long been discussed in the literature (Marie et al., 2005; Shukla et al., 2007; Kim et al., 2015; Kosty and Vogel, 2015; Pfaff et al., 2016; Timberlake et al., 2017). However, our top hit gene-set is KEGG’s cancer pathway, and this link between CRS and cancer is to our knowledge rarely discussed in the literature. In Table 3, we list few more cancer pathways, though not family-wide significant, that are significant at *p* = 10^−5^ level.

**Table 3:** Cancer-related gene-sets in MSigDB C2 that just missed the family-wide significance level, and the corresponding overlapping gene names. Gene-set size, overlapping set size, Fisher’s test p-value and odds-ratio (OR) are also listed.

| potentially interesting enrichment for cancer-related gene-sets in C2 |  |  |  |  |
| --- | --- | --- | --- | --- |
| gene-set name | gs-size | match | fisher-pv | OR |
| **colorectal cancer (KEGG)**<br>AXIN2, BRAF, KRAS, RAF1, TGFBR1, TGFBR2 | 62 | 6 | 4.1E-6 | 16.2 |
| **pancreatic cancer (KEGG)**<br>BRAF, KRAS, RAF1, STAT3, TGFBR1, TGFBR2 | 70 | 6 | 7.8E-6 | 14.4 |
| **Greshock cancer copy number**<br>BRAF, FGFR1, FGFR2, FGFR3, GNAS, KAT6B<br>KRAS, PTPN11, RECQL4, SMO, TCF12 | 322 | 11 | 7.4E-6 | 5.9 |
| **melanoma (KEGG)**<br>BRAF, FGF9, FGFR1, IGF1R, KRAS, RAF1 | 71 | 6 | 8.5E-6 | 14.2 |
| **CML (KEGG)**<br>BRAF, KRAS, PTPN11, RAF1, TGFBR1, TGFBR2 | 73 | 6 | 9.8E-6 | 13.8 |

The C3 collection in MSigDB defines gene-sets by their shared DNA sequence motif in the promoter region. With a p-value threshold of 5.7 × 10^−7^ and 1.2 × 10^−5^ (0.01/836), 1 or 6 gene-sets are family-wise significant. Three of our six enriched gene-sets (AACTTT, TTGTTT, CTTTGT) appear in the top ten enriched C3 gene-sets in SOX10 regulated genes (supplementary Table 5 of (Sun et al., 2014)). Similar to SOX6 (Hagiwara, 2011), one of the CRS genes, SOX10 also plays an important role in embryonic development (Honoré et al., 2003). These results and matching genes are listed in Table 4, and volcano plot in Fig.1(C).

**Table 4:** Gene-sets in MSigDB C3 (motif) that are enriched with craniosynostosis genes (significant at the family-wide level), and the corresponding overlapping gene names. Gene-set size, overlapping set size(m), Fisher’s test p-value (pv) and odds-ratio (OR) are also listed.

| C3 (motif) |  |  |  |  |
| --- | --- | --- | --- | --- |
| gene-set: motif (TF or micro RNA) | gs-size | m | pv | OR |
| **AACTTT (IRF1?)**<br>ALX4, AXIN2, BCOR, COLEC10, CYP26B1, EFNA4, ERF, FGF9, FGFR1, FGFR2, GNAS, GPC3, IGF1R, IL6ST, IRAX5, JAG1, KIAA1267, KAT6A, MLL2, KRAS, LMNA, MSX2, NFIA, PHEX, PTH2R, RAB23, RSPRY1, SMAD6, SOX6, SPRY1, STAT3, TCF12, ZEB2, ZIC1, ZNF462 | 1820 | 35 | 1E-9 | 4 |
| **ATAAGCT (mir21)**<br>HNRNPK, JAG1, PPP3CA, SKI, SOX6, SPRY1, STAT3, TGFBR2 | 112 | 8 | 8.3E-7 | 12.2 |
| **TTGTTT (FOXO4.01)**<br>ASXL1, AXIN2, BCOR, CD96, CTSK, DPH1, EFNA4, EFNB1, FGF9, GNAS, GPC3, IGF1R, IL6ST, JAG1, LRP5, MASP1, MSX2, NFIA, NTRK2, PHEX, PPP1CB, PTPRD, RAB23, RUNX2, SHOC2, SMAD6, SOX6, STAT3, TWIST1, ZEB2, ZIC1 | 1979 | 31 | 1.2E-6 | 3.1 |
| **AAAGGGA (mir204/mir211)**<br>ALPL, FREM1, GLIS3, MED13L, NTRK2, P4HB, RUNX2, SEC24D, TCF12, TGFBR2 | 219 | 10 | 1.7E-6 | 7.9 |
| **CTTTGT (LEF1.Q2)**<br>ATR, AXIN2, BRAF, CD96, EFNA4, EFNB1, FAM20C, FGF9, FGFR3, GLIS3, GPC3, HNRNPK, IGF1R, IRX5, KAT6A, MLL2, KRAS, LRP5, MASP1, PPP1CB, RSPRY1, SMAD6, SOX6, STAT3, TCF12, TCOF1, TGFBR1, ZEB2, ZIC1 | 1892 | 29 | 4.5E-6 | 2.9 |
| **RYTTCCTG (ETS2.B)**<br>BMP2, BRAF, CD96, COLEC11, CTSK, ERF, FGFR2, HNRNPK, IFT43, IL11RA, KIAA1267, MLL2, LMX1B, LRP5, NFIA, PHEX, PTPN11, TCF12, TGFBR2, ZEB2 | 1048 | 20 | 8.9E-6 | 3.5 |

Although AACTTT/AAAGTT is not labeled as a binding motif for a known transcription factor (TF), IRF1 is listed as the binding TF for this motif in (Xie et al., 2005). Among other binding partners, mir21 (Feng and Tsao, 2016; Javanmardi et al., 2017), mir204 (Li et al., 2016), mir211 Mazar et al., (2016), FOXO4 (Liu et al., 2011; Hornsveld et al., 2018), LEF1 (Hovanes et al., 2001; Li et al., 2009), ETS2 (Sizemore et al., 2017; Fry and Inoue, 2018) are all discussed in the context of a cancer.

The C5 collection in MSigDB is defined by the Gene Ontology (GO) terms (Gene Ontology Consortium, 2019). With the number of all MSigDB gene-sets to correct multiple testing, 111 GO gene-sets are significant; with only the number of GO gene-sets (*n*_*GS*_ = 5917) to correct, 126 GO gene-sets are significant. We highlight two groups of significant GO gene-sets: the first group has higher percentage of genes in the overlapping set (Table 5, for percentage > 0.1), and the second group contains larger gene-sets (Table 6, for gene-set size > 1000 genes). These two groups as marked in the volcano plot (Li, 2012) can be seen at Fig.1(C). The larger-sized gene-sets are located near the inner side of the volcano as these are “common events” (Li et al., 2014), whereas the stronger-signal gene-sets are located at the outer side because these are relatively rare events.

**Table 5:** Partial gene-sets in MSigDB C5 (gene-ontology) that are enriched with craniosynostosis genes (significant at the family-wide level), with number of overlapping genes (m) at least 10% of the gene-set size. Fisher’s test p-value (pv) and odds-ratio (OR) are also listed. act: activity; dev: development; diff: differentiation; mem: membrane; morphog: morphogenesis; neg: negative; pos: positive; prot: protein; rec: receptor; reg: regulation; sig: signaling; sys: system;

| C5 (gene ontology), stronger signal (10% genes of gene-set overlap) |  |  |  |  |
| --- | --- | --- | --- | --- |
| gene-set name | gs-size | m | pv | OR |
| bone dev | 155 | 22 | 1E-22 | 27.8 |
| skeletal sys morphog | 200 | 23 | 8E-22 | 22.8 |
| bone morphog | 79 | 16 | 6.3E-19 | 37.5 |
| cranial ske dev | 55 | 14 | 6.7E-18 | 46.2 |
| reg ossification/pos/neg | 174/82/67 | 19/9/7 | 9.3E-18/5.2E-9/3.8E-7 | 21.1/18.9/17.7 |
| appendage dev | 166 | 18 | 8.1E-17 | 20.8 |
| reg osteoblast diff/pos | 109/58 | 15/9 | 1.4E-15/3.2E-10 | 25.2/26.8 |
| reg smoothened sig/pos | 62/24 | 11/5 | 8.6E-13/9E-7 | 31.3/34.7 |
| embryonic skeletal sys morphog | 93 | 12 | 2.3E-12 | 22.9 |
| embryonic skeletal sys dev | 122 | 13 | 2.4E-12 | 19.1 |
| odontogenesis | 105 | 12 | 8.3E-12 | 20.9 |
| embryonic cranial skeleton morphog | 46 | 9 | 5.1E-11 | 33.8 |
| *smad binding* | 70 | 10 | 6.7E-11 | 24.9 |
| biomineral tissue dev | 75 | 10 | 1.2E-10 | 23.2 |
| chondrocyte diff | 60 | 9 | 4.2E-10 | 25.9 |
| reg cartilage dev/pos | 62/29 | 9/6 | 5.5E-10/7.2E-8 | 25.1/34.8 |
| craniofacial suture morphog | 14 | 6 | 1.8E-9 | 75.2 |
| reg chondrocyte diff/pos | 45/19 | 8/5 | 1.2E-9/3.3E-7 | 30.4/43.9 |
| *trans-mem rec prot kinase/tyrosine* | 81/64 | 9/7 | 4.7E-9/2.9E-7 | 19.2/18.5 |
| bone mineralization | 38 | 7 | 1.1E-8 | 31.3 |
| ciliary tip | 43 | 7 | 2.4E-8 | 27.6 |
| **reg mesenchymal cell proliferation/pos** | 34/27 | 6/6 | 1.7E-7/4.9E-8 | 29.7/39.6 |
| embryonic digit morphog | 59 | 7 | 1.7E-7 | 20.1 |
| **pos reg stem cell proliferation** | 61 | 7 | 2.1E-7 | 19.5 |
| body morphog | 44 | 6 | 6.5E-7 | 22.9 |
| endochondral bone morphog | 45 | 6 | 7.3E-7 | 22.4 |
| *prot transport along microtubule* | 27 | 5 | 1.5E-6 | 30.9 |

**Table 6:** Partial gene-sets in MSigDB C5 (gene-ontology) that are enriched with craniosynostosis genes (significant at the family-wide level), with at least 1000 genes in the gene-set. Number of overlapping genes (m), Fisher’s test p-value (pv) and odds-ratio (OR) are also listed. dev: development; diff: differentiation; morphog: morphogenesis; neg: negative; org: organismal; pos: positive; proc: process; prot: protein; prom: promoter; reg: regulation; res: response; sys: system; tran: transcription;

| C5: gene ontology, large gene-set size (> 1000 genes); or very significant |  |  |  |  |
| --- | --- | --- | --- | --- |
| gene-set name | gs-size | match | fisher-pv | OR |
| tissue dev | 1501 | 43 | 4.7E-18 | 6.8 |
| reg cell diff | 1470 | 41 | 9.6E-17 | 6.4 |
| reg multicell org dev | 1652 | 39 | 1.5E-13 | 5.2 |
| pos reg dev | 1129 | 32 | 3.9E-13 | 5.9 |
| reg tran (RNA polII prom) | 1763 | 39 | 1.1E-12 | 4.9 |
| *res endogenous stimulus/cellular..* | 1440/1000 | 35/25 | 1.8E-12/31.E-9 | 5.2/4.8 |
| **reg cell proliferation** | 1478 | 35 | 3.7E-12 | 5 |
| cell res org substance | 1833 | 38 | 1.7E-11 | 4.5 |
| pos reg biosynthetic proc | 1792 | 36 | 1.7E-10 | 4.2 |
| pos reg gene expression | 1719 | 35 | 2.3E-10 | 4.3 |
| cell dev | 1406 | 31 | 5.1E-10 | 4.5 |
| *pos reg resp stimulus* | 1878 | 36 | 5.9E-10 | 4 |
| pos reg multicell org proc | 1382 | 30 | 1.5E-9 | 4.4 |
| neurogenesis | 1388 | 30 | 1.7E-9 | 4.3 |
| *pos/neg reg cell communication* | 1520/1176 | 31/27 | 3.3E-9/3.8E-9 | 4.1/4.5 |
| reg anatomical struture morphog | 1011 | 25 | 3.8E-9 | 4.8 |
| reg cell death | 1458 | 29 | 2.1E-8 | 3.9 |
| immune sys proc | 1927 | 32 | 2.1E-7 | 3.3 |
| skeletal sys dev | 453 | 37 | 3.1E-30 | 18.9 |
| org morphog | 839 | 42 | 1.8E-26 | 12.1 |

Most family-wide significant GO gene-sets are expected to be related to CRS as they are involved in morphogenesis, development, differentiation, ossification. Once again, we see some significant gene-sets to be related to cell proliferation. A few significantly CRS-associated gene-sets are related to protein binding, protein transport, signal transduction, cell communication, etc.

### Enrichment analysis of craniosynostosis genes by Ingenuity pathway analysis

We ran an enrichment analysis of 113 CRS genes by the Ingenuity Pathway Analysis (IPA) (using the Ingenuity Knowledge Base). The top 16 canonical pathways are shown in Fig.2, with the p-value cutoff of 1.32E-5. In Fig.2, signaling pathways are marked by blue, and cancer gene-sets are marked by red, with the rest being well known to be related to development and differentiation. The presence of the “molecular mechanisms of cancer” gene-set in the top list once again confirms the link between CRS and cancer.

**Figure 2:**
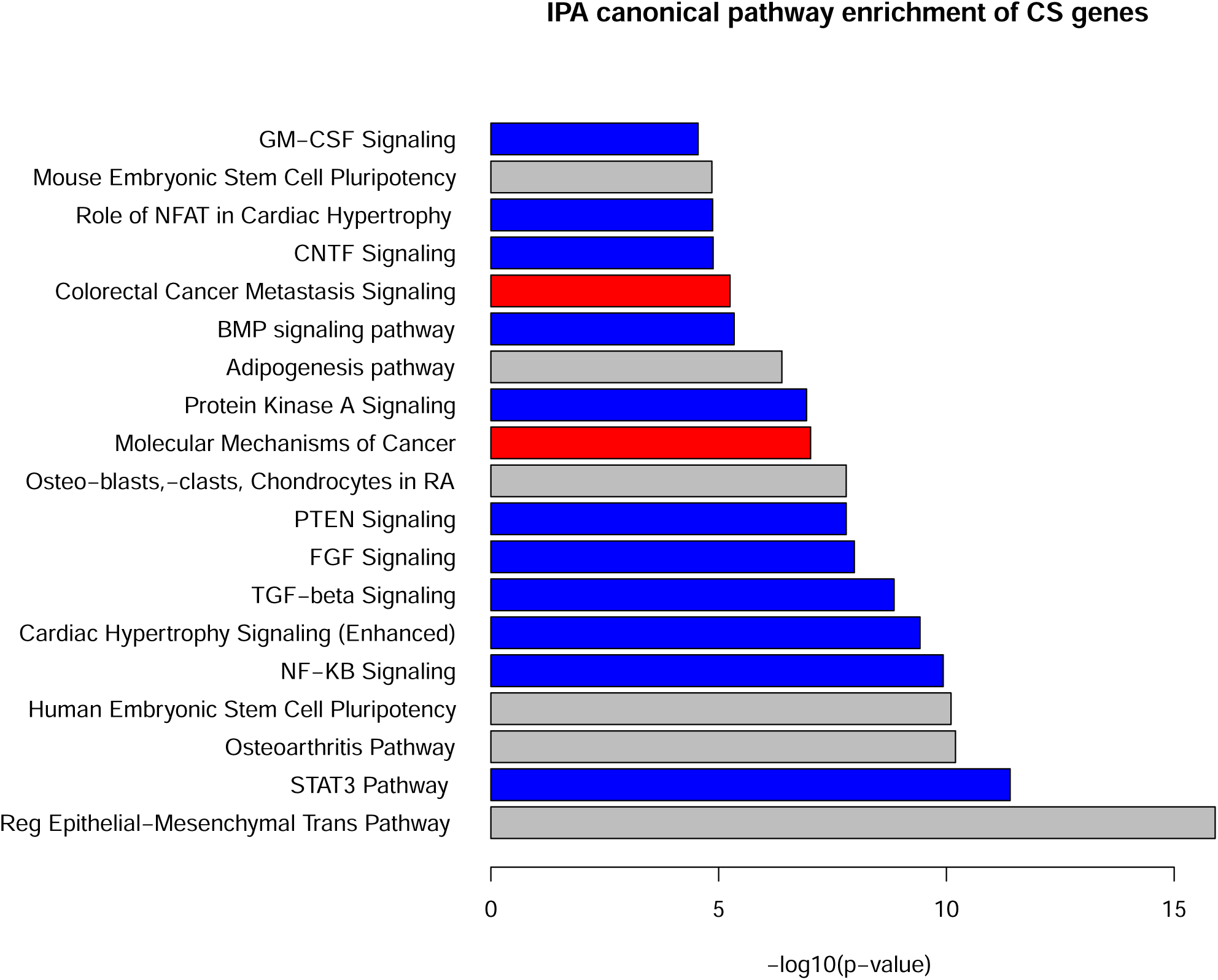
IPA canonical pathways that are significantly enriched with craniosynostosis genes at p-value of 3×10^−5^ level. The cancer related pathways are colored by red, and signaling pathways by blue. The remaining pathways are all related to development and differentiation.

It is interesting that some IPA cancer genes are not in KEGG cancer pathway list or cancer driver gene list, which lead to some difference between the overlapping results. More specifically, of the three overlapping lists (*n*_1_ = 21 for driver genes, *n*_2_ = 15 for KEGG cancer genes, and *n*_3_ = 13 for IPA molecular mechanism of cancer genes), only four genes appear in all three lists: BRAF, KRAS, RAF1, and TGFBR2. Besides these four genes, there are also pairwise shared genes: AXIN2, FGFR1, FGFR2, and FGFR3 between the driver and KEGG lists, BMP2, SMO, and TGFBR1 between the driver and IPA lists, and ATR, GNAS, and PTPN11 between the KEGG and IPA lists. The remaining genes are unique to specific lists: ASXL1, BCOR, FLNA, HUWE1, IL6ST, KANSL1, KMT2D, PTPRD, SMCIA, and TCF12 for the driver list, FGF9, GLI3, IGF1R, and STAT3 for the KEGG list, and IHH, LRP5, and SMAD6 for the IPA list.

More detailed information of those IPA signaling pathways in Fig.2 are included in Table 7. Among signaling pathways, IPA shares some common results with KEGG/Reactome/MSigDB, such as TGF and FGF, though there are other significant hits, such as STAT3, NF-KB, PTEN, etc. The decision to put certain genes in a pathway, and the definition of a pathway in light of cross-talk among pathways (Attisano and Labbé, 2004; Guo and Wang, 2009; Mendoza et al., 2011), may all contribute to these discrepancies. It justified our use of multiple databases for this analysis.

**Table 7:**
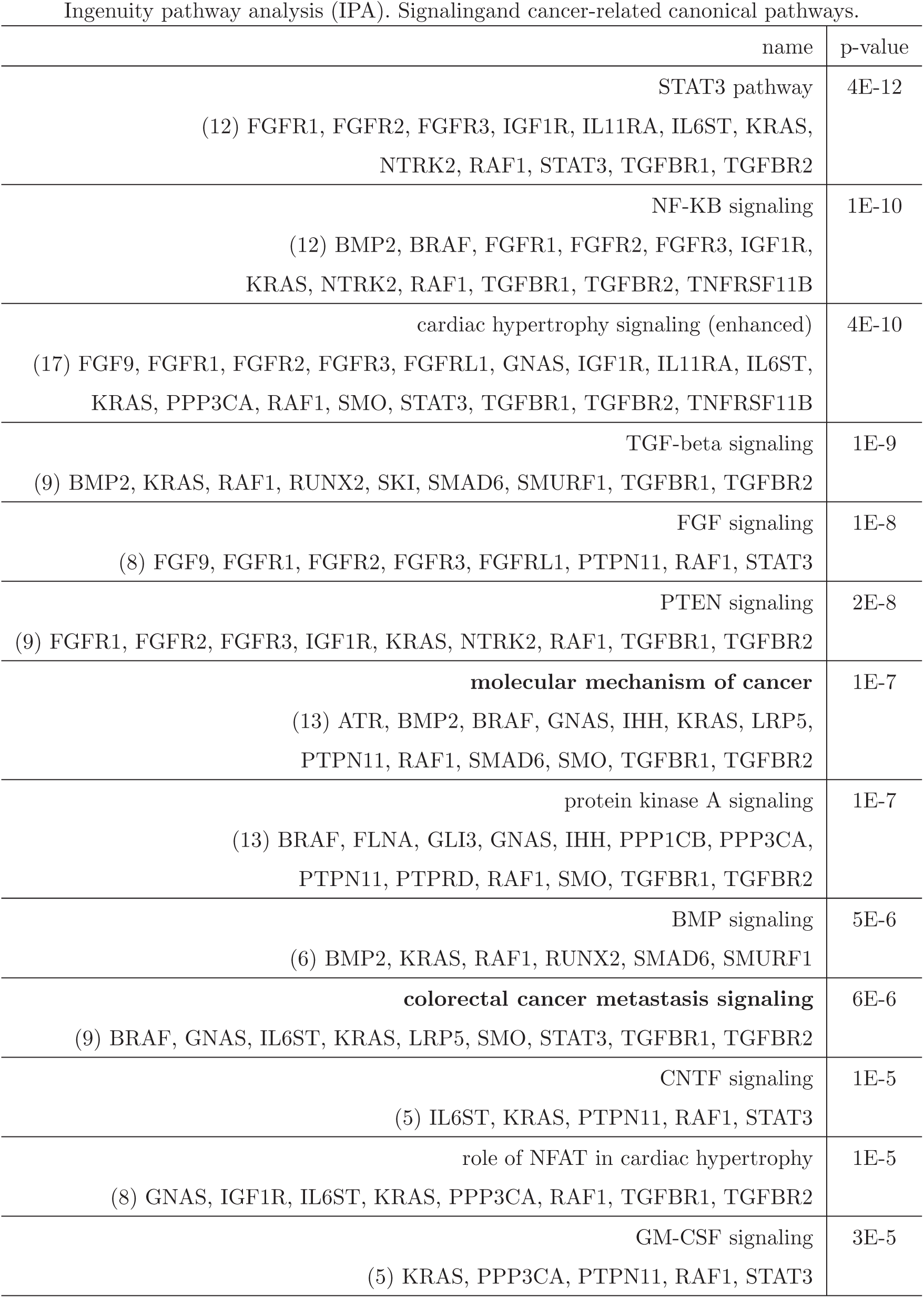
More information on the IPA canonical pathways in Fig.2 (enrichment p-value < 3 × 10^−5^) that are either a signaling pathway, or related to cancer: number and names overlapping genes and enrichment p-value.

Besides the IPA canonical pathways, Table 8 shows top-5 hits in other categories: upstream regulators, diseases, molecular/cellular functions, physiological system development and function. Most of these hits are expected due to our knowledge of CRS being a developmental disorder. We do notice cell-signaling as one of the top hits in molecular/cellular functions.

**Table 8:**
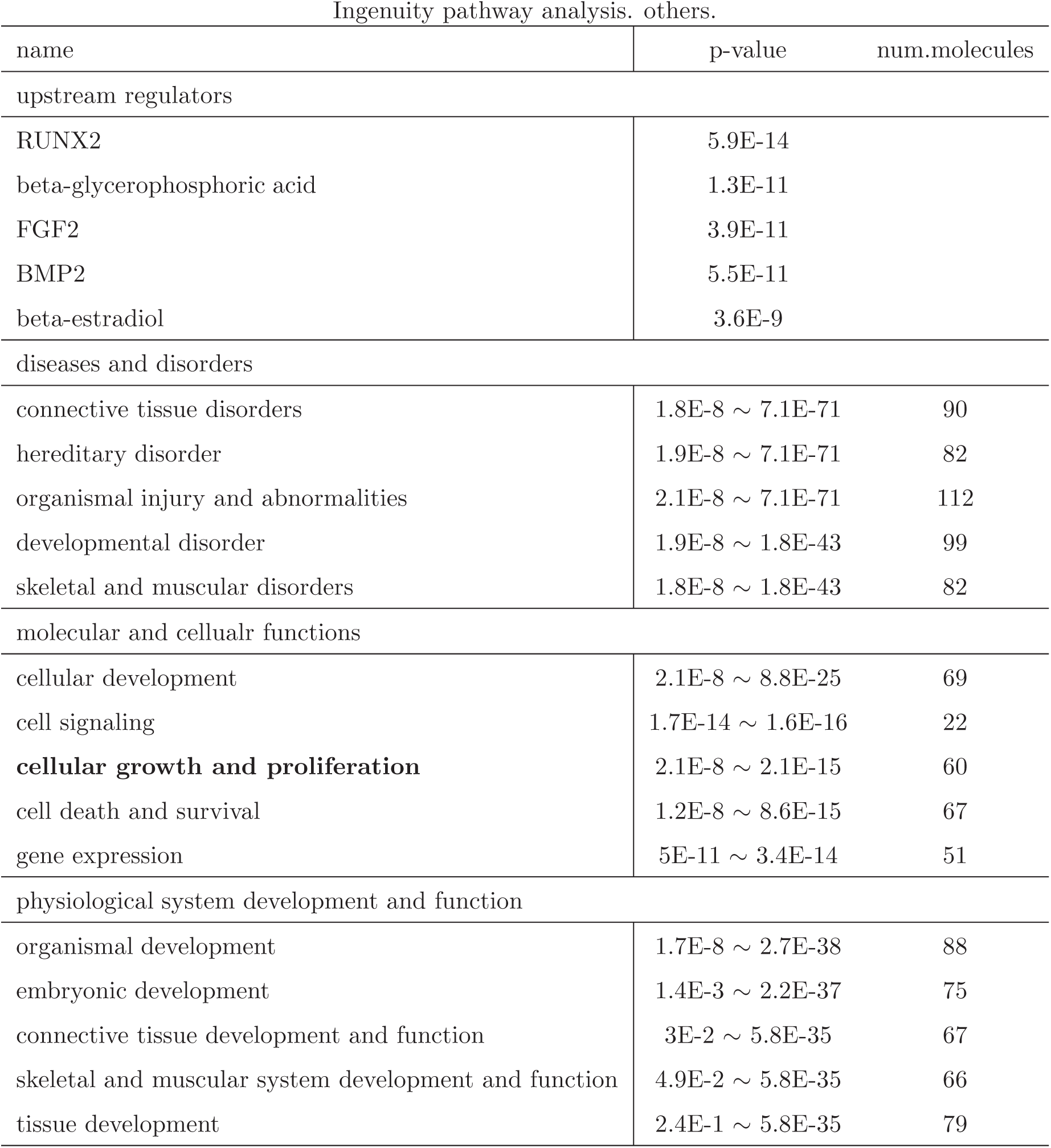
IPA top five hits in other categories, based on enrichment/association analysis with the craniosynostosis genes: upstream regulators, diseases and disorders, molecular and cellular functions, physiological system development and function.

In order to find out which specific CRS genes participate in which signaling pathway, we checked the 15 CRS genes that overlap with the KEGG cancer pathway. Their presence in other MSigDB C2 signaling pathways and IPA signaling pathways is summarized in Table 9. From Table 9, a few “core” genes appear in multiple IPA signaling pathways (FGFR1, FGFR2, FGFR3, KRAS, RAF1, STAT3, TGFBR1, TGFBR2), which other are more specific for one or few signaling pathways. For example, SMO only appears in cardiac hypertrophy and protein kinase A signaling pathways; GLI3 only appears in protein kinase A signaling pathways; BRAF only appears in NF-KB and protein kinase A signaling pathways; etc.

**Table 9:**
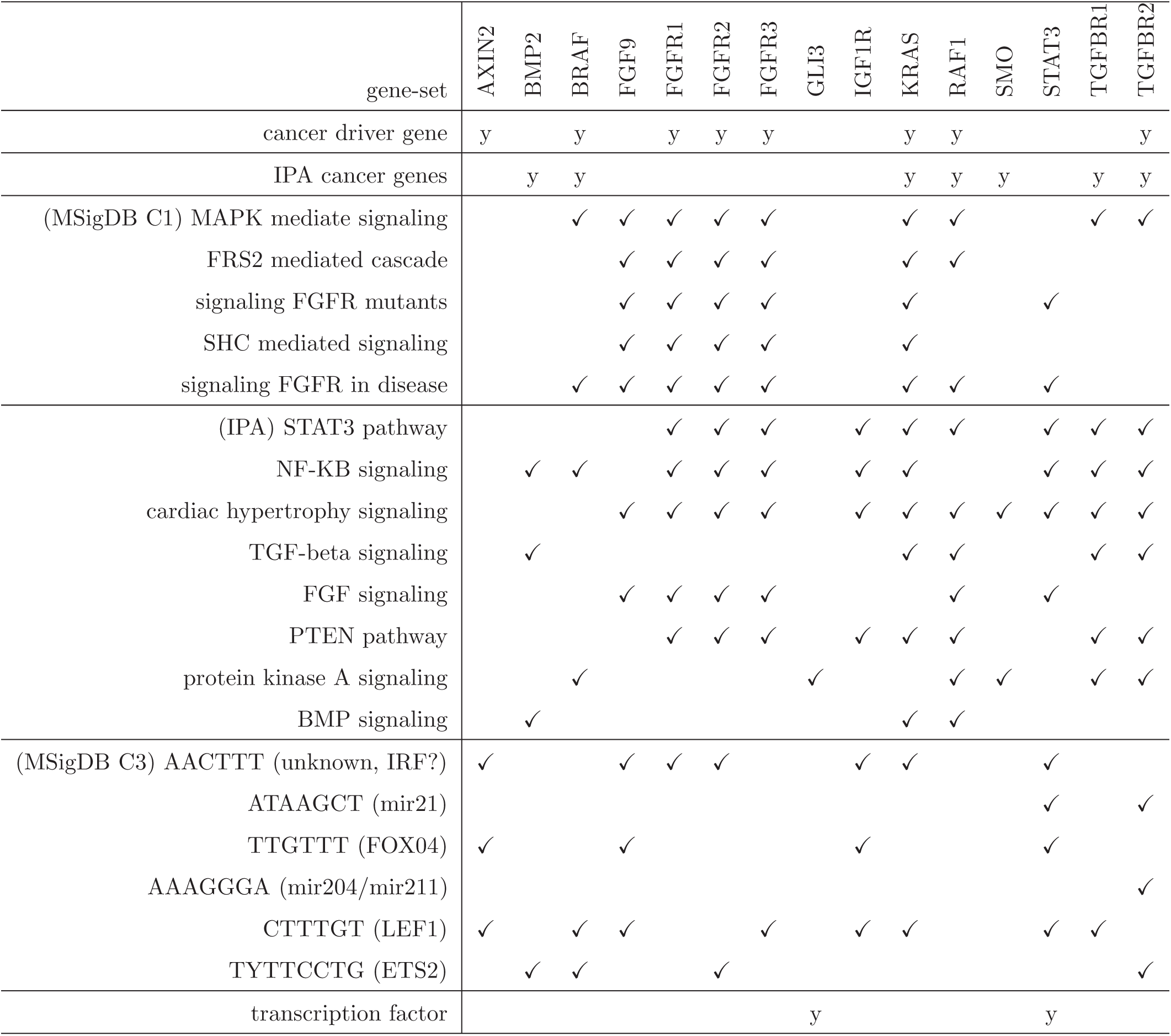
Fifteen craniosynostosis genes that are also in KEGG cancer pathway: their presence in cancer driver gene list; in IPA cancer gene list; in other MSigDB C2, C3 gene-sets; in IPA signaling pathways; and in a transcription factor list (Lambert et al., 2018).

### Enrichment analysis of craniosynostosis genes in transcription factors

Transcription factors (TF) can be the end target of a signaling pathway. Due to the enrichment of signaling pathway genes in CRS gene list, we wonder whether TFs are enriched in the list also. There have been previous attempts to examine the connection between TF and diseases (Latchman, 1996; Li et al., 2015a), and it would not be completely unexpected that the CRS gene list contains a higher proportion of TFs. MSigDB does not have a gene-set with a complete list of transcription factors. The closest ones include the geneontology set in C5 “GO_SEQUENCE_SPECIFIC_DNA_BINDING” with around 1000 genes, and “GO_REGULATION_OF_TRANSCRIPTION_FROM_RNA_POLYMERASE_II_PROMOTER” with around 1700 genes.

Instead, we used a recently compiled TF list from (Lambert et al., 2018) with 1638 TFs. There are 19 (out of 113, or 16.8%) CRS genes present in the TF list, compared to the baseline TF frequency of 8.2% (1638/20000). The Fisher test of the enrichment has p-value = 0.003. These overlapping genes are: AHDC1, ALX4, ERF, GLI3, GLIS3, IRX5, LMX1B, MSX2, NFIA, RUNX2, SIX2, SKI, SOX6, STAT3, TCF12, TWIST1, ZEB2, ZIC1, and ZNF462.

There is another TF gene list from ten years ago with 1835 TFs (Vaquerizas et al., 2009). The overlap between the old and new TF lists is 1423 genes (87% of the new list, and 78% of the old list). Some CRS genes that are present in the old TF list, e.g., BMP2, SMAD6, are said to not be DNA binding proteins with specific sequence motifs (Lambert et al., 2018). Though we may trust less the TF list from ten years ago than the recent list, if we use the old TF list for enrichment test, it contains 22 CRS genes (22/113=16.5%), vs. the baseline frequency of 9.2% (1835/20000), with a Fisher test p-value of 0.0008.

## Methods and Data

### Statistical analysis

The Fisher’s test for 2-by-2 table is by the R (*www.r-project.org*) function *fisher.test*. Other data manipulation, text parsing, and data plotting tasks are carried out by our own scripts in perl, python, and R.

### Gene intolerance databases

Three databases on whether a gene is tolerant to functional mutation are used: (1) Residual Variation Intolerance Score (RVIS): *genic-intolerance.org*. The RVIS values are based on ExAC (Exome Aggregation Consortium) v2 (release 2.0). (2) Loss-of-function Observed/Expected Upper bound Fraction (LOEUF): *gnomad.broadinstitute.org/downloads*, follow the link of “Constraint”, then “pLoF metrics by gene TSV”. The gene-level metrics are determined from the gnomAD version 2.1.1 (Karczewski et al., 2019). (3) Gene Damage Index (GDI): *lab.rockefeller.edu/casanova/GDI*. The GDI values are based on the healthy individual’s whole genome sequence data from the 1000 Genomes Project database (phase 3) (The 1000 Genomes Project Consortium, 2015), and the impact/damage from a variant is determined by the CADD method (Kircher et al., 2014).

### Gene-set database

We downloaded the gene-set data from MSigDB (*software.broadinstitute.org/gsea/msigdb/collections.jsp*), version 6.2 (Liberzon et al., 2015). There are 17810 gene-sets in total, and we excluded the C1 collection which contains chromosome location based gene-sets, leaving 17484 gene-sets. We also filter out non-coding genes, according to the annotation from HGNC (*www.genenames.org*), from the gene-sets. The remaining gene-set collections used are: H (Hallmark collection, *n*_*GS*_=50), C2 (curated gene-sets, including KEGG (*www.genome.jp/kegg*), REACTOME (*reactome.org*), BioCarta, Canonical Pathways, Chemical and Genetic Perturbation, *n*_*GS*_=4762), C3 (motif gene-sets, n=836), C4 (cancer gene neighborhoods (Subramanian et al., 2005) and cancer modules (Segal et al., 2004), *n*_*GS*_=858), C5 (Gene Ontology (*geneontology.org*) gene, *n*_*GS*_=5917), C6 (GEO microarray derived cancer signatures *n*_*GS*_=189), and C7 (immunologic signatures (Godic et al., 2016), *n*_*GS*_=4872).

### Cancer driver gene lists

The 2018 list with 299 cancer driver genes is downloaded from Supplement Table S1: *https://www.sciencedirect.com/science/article/pii/S009286741830237X#app2* of the paper (Bailey et al., 2018) (doi: 10.1016/j.cell.2018.02.060). The 2013 list with 125 driver genes is downloaded from the Supplementary Table S2A: *www.sciencemag.org/cgi/content/full/339/6127/1546/DC1* in the paper (Vogelstein et al., 2013) (doi: 10.1126/science.1235122).

### Ingenuity pathways

Ingenuity Systems is a commercial package (©2000-2019 QIAGEN) with an expert-curated Knowledge Base, including canonical pathways.

### Transcription factor lists

The 2018 list of 2764 transcription factors is downloaded from *humantfs.ccbr.utoronto.ca/*, also as a supplement Table S1 in (Lambert et al., 2018) (doi: 10.1016/j.cell.2018.01.029). The 2009 list of transcription factors is downloaded from the supplementary Table 2 of (Vaquerizas et al., 2009) (doi: 10.1038/nrg2538).

## Discussion

### The statistical evidence linking craniosynostosis and cancer is robust

We have obtained strong statistical evidence (p-value ∼ 10^−16^) that craniosynostosis genes and cancer driver genes sets are more overlapping than by chance. The evidence is even more convincing that when craniosynostosis genes list is split into three pieces according to the time when it was compiled, there is an over-representation for each sub-list. Furthermore, even when we use an older cancer driver gene list with half of the genes as the new list, thus smaller sample sizes, the enrichment test is still significant (p-value ∼ 10^−7^). The craniosynostosis-cancer connection is further confirmed in two other databases: the KEGG cancer pathway and the IPA molecular mechanism of cancer gene-set, despite the difference between these cancer gene lists.

### Redundancy between gene-sets is not enough to carry over an enrichment between craniosynostosis gene list and one gene-set and that of another

It is known that gene-set databases contain redundant gene-sets (Vivar et al., 2013; Stoney et al., 2018). In order to check whether redundancy is the reason that we obtained multiple enrichment gene-sets in Table 2, we examine the self-enrichment between KEGG_PATHWAYS_IN_CANCER and other C2 and C3 MSigDB gene-sets. Fig.3 shows enrichment Fisher’s test (− log_10_) p-value as a function of the C2/C3 gene-set sizes. The points (C2/C3 gene-sets) at the top of the plot exhibit significant overlap with the KEGG cancer pathway gene-set. The gene-sets that significantly overlap with craniosynostosis gene list are marked in red.

**Figure 3:**
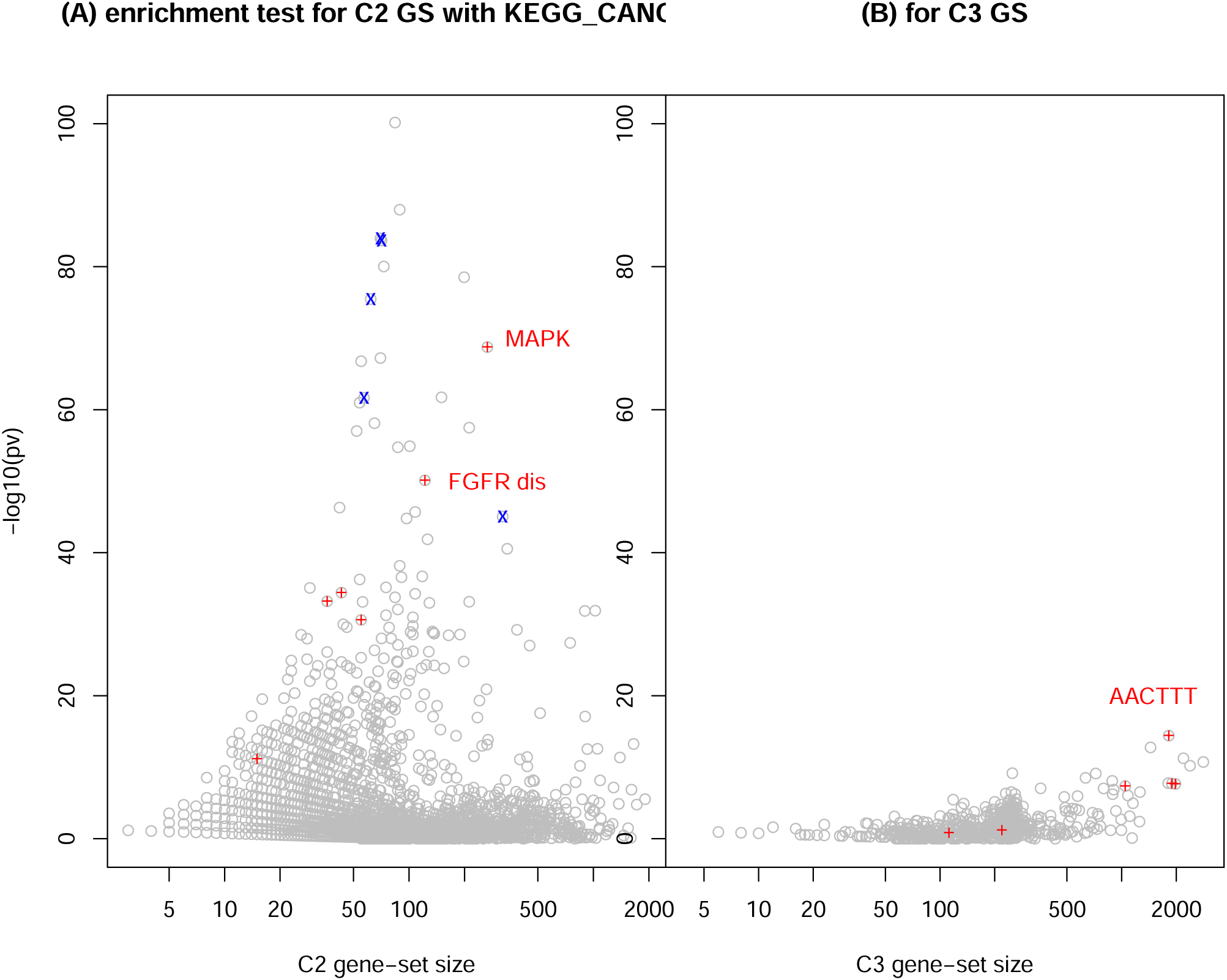
Enrichment of KEGG cancer pathway genes in other MSigDB C2 and C3 gene-sets. The *x*-axis is the the gene-set size (in log-scale), and the *y*-axis is the minus log10 of Fisher’s test p-value. In (A), the red dots are C2 gene-sets listed in Table 2, and blue dots are the cancer-related C2 gene-sets listed in Table 3. In (B), the red dots are C3 gene-sets listed in Table 4.

Indeed, KEGG_MAPK_SIGNALING_PATHWAY shares 82 out of 266 in common with the KEGG cancer gene-set, REACTOME_SIGNALING_BY_FGFR_IN_DISEASE shares 53 out of 122 in common, followed by SIGNALING_BY_FGFR (Reactome): 31/43, FRS2_MEDIATED_CASCADE (Reactome): 29/36, FGF (PID): 30/55, and SHC_MEDIATED_SIGNALING (Reactome): 10/15. For C3 gene-sets, AACTTT motif defined gene-set shares 83/1820 with the KEGG cancer list, once again, indicating a potential link with the cancer. The C2 cancer gene-sets listed in Table 3 are marked in blue in Fig.3, which are all highly correlated with the KEGG cancer gene-set. However, a high enrichment with the KEGG cancer gene-set does not guarantee an enrichment with the craniosynostosis gene list. For example, the top two points in Fig.3(A) are KEGG small-cell lung cancer gene-set and KEGG prostate cancer gene-set. While the prostate cancer gene-set has some overlap with craniosynostosis gene list (6 out of 89, p-value=3×10^−5^), the small-cell lung cancer gene-set actually has zero overlap with the craniosynostosis gene list.

### Signaling pathways as the main cause for craniosynostosis-cancer connection

Our results in Tables 2,7 and Fig.3(A) all indicate the importance of signaling pathways in craniosynostosis, as well as their common link to both craniosynostosis and cancer. This is consistent with previous findings. FGFR2 is the first gene highlighted to be involved in both craniosynostosis and cancer (Pollock et al., 2007). The FGF signaling pathway has been well established to participate both morphogenesis/development/differentiation (Deimling and Drysdale, 2011; Drewer et al. 2016), on one hand, and carcinogenesis, on the other (Ahmed et al., 2012; Fearon et al., 2013; Gallo et al., 2016). The FGF signaling pathway was proposed as a drug target for treating cancers (Lee and Secord, 2014; Porta et al., 2017; Hanrahan et al., 2019). The WNT signaling pathway is associated with craniosynostosis (Timberlake et al., 2016), and is also associated with cancer (Clevers, 2006; Clevers and Nusse, 2012; Zhan et al., 2017). The Hedgehog signaling pathway is discussed in connection to craniosynostosis (Jenkins et al., 2007), and is also associated with cancer (Jiang and Hui, 2008; Skoda et al., 2018).

In Table 9 we can see that all craniosynostosis-KEGG-cancer overlapping genes are components of some signaling pathways (not mentioned in Table 9, AXIN2 is part of the WNT signaling pathway, BMP2, GLI3, and SMO are part of the Hedgehog signaling pathway, IGF1R’s binding partner, IGF1, is in the P53, mTOR, and PIP3 signaling pathways). Among 16 more craniosynostosis genes that are not in Table 9 but in the cancer driver list or in the IPA molecular mechanisms of cancer list, FLNA, GNAS, IHH, IL6ST, LRP5, PTPN11, PTPRD, and SMAD6 are also in one of the significantly enriched signaling pathways in Tables 2 and 7. The existence of so many three-way overlapping genes (craniosynostosis, cancer, signaling) is evidence that signaling pathways might be behind the craniosynostosis-cancer connection.

### Transcription factors and microRNA associated with craniosynostosis

A biological process can be traced from the chemical signal sent from outside the cell, through the cell membrane, through intra-cellular signaling pathways, into the nucleus, to trigger transcription factors, bind to promoters of certain genes, and initiate transcription. From this description, both transcription factors and their targets can be involved in the etiology of a disease. We have already seen that TFs are enriched in craniosynostosis genes. We can also ask the same question on targets of TFs.

Table 4 shows six TF-motif defined gene-sets are enriched in the craniosynostosis gene list. It hints, though does not prove, that FOXO4, LEF1, ETS2, and possibly IRF1, are relevant to craniosynostosis. A similar point can also be made on micro RNA regulation: mir21 and mir204/mar211. Some craniosynostosis-cancer overlapping genes do not appear in any signal pathways, but they are present in some motif-defined significantly enriched gene-sets, e.g. ASXL1, BCOR, TCF12. These results may guide us to investigate further their role in both craniosynostosis and in cancer.

Interestingly, TF genes are also enriched in cancer driver genes. Using the new TF list from (Lambert et al., 2018), 51 out of 299 cancer driver genes from (Bailey et al., 2018) (17%) are TFs, as compared to the baseline TF frequency of 8%. The Fisher’s test on enrichment has p-value = 7.8×10^−7^. The participation of TF in tumorigenesis is widely studied (Darnell, 2002; Harris, 2002; Zheng and Blobel, 2011; Gupta et al., 2019), including the most famous TF p53 (TP53 gene) (Harris, 1993; Staib et al., 2003; Muller and Vousden, 2013). However, when the three gene lists are compared (TF, cancer driver, CRS), only one gene, TCF12 (Lee et al., 2012; Sharma et al., 2013), appears in all lists.

### The oncogenesis recapitulating ontogenesis framework

Julius Cohnheim (1839-1884) was perhaps the first person to propose the embryonic origin of cancer (Capp, 2019), influenced by his teacher Rudolf Virchow (1821-1902) (Ackerknecht, 1958). This parallel between normal developmental process in embryo and cancer in adults can be called “oncogenesis recapitulating ontogenesis” (Huang et al., 2009). Note that there is also a perceived parallel between individual development and species evolution, called “ontogeny recapitulating phylogeny” (Swinnerton, 1938; Richardson and Keuck, 2002).

There are multiple similarities between development and cancer, nicely summarized in (Naxerova et al., 2009). The microscope image of some cancer tissues look like undifferentiated immature cells (Rosai et al., 1985). Cell migration during embryonic development is similar to cancer metastasis (Ridley et al., 2003). More recently, expression profiling (transcriptome) of both certain cancer and developmental tissues exhibit similar signatures (Borczuk et al., 2003; Kho et al., 2004; Coulouarn et al., 2005; Hu and Shivdasani, 2005; Liu, 2006; Dekel et al. 2006; Kaiser et al., 2007; Naxerova et al., 2009; Li et al., 2010; Palmer et al., 2012; Soundararajan et al., 2015). There are also many theoretical hypotheses and frameworks building around the idea (Garraway and Sellers, 2006; Ao et al., 2008; Huang et al., 2009; Huang, 2011; Yuan et al., 2017; Zhou et al., 2018; Liu, 2018, 2019) The results in our paper add another path linking development and cancer.

In conclusion, we provide solid evidence that a higher proportion of genes associated with craniosynostosis are also linked to cancer. This observation is consistent with the perspective that adult carcinogenesis (oncogenesis, tumorigenesis) and embryonic ontogenesis (morphogenesis) share certain biological process in common, such as signaling pathways. Transcription factors, as end points of signaling pathways, also play prominent roles in both craniosynostosis and cancers.

## abbreviations

CADD: combined annotation dependent depletion;
CML: chronic myelogenous leukemia;
CNV: copy number variation;
CRS: craniosynostosis;
FGF: fibroblast growth factor;
GDI: gene damage index;
GEO: gene expression omnibus database;
GO: gene ontology;
GS: gene-set;
GWAS: genome-wide association studies;
HUGO: Human Genome Organisation;
IPA: Ingenuity Pathway Analysis;
KEGG: Kyoto Encyclopedia of Genes and Genomes;
LOEUF: loss-of-function observed-over-expected upper bound fraction;
MeSH: medical subject headings;
MSigDB: Molecular Signature Database;
OR: odds-ratio;
PID: pathway interaction database;
pv: p-value of a test;
RVIS: residual variation intolerance score;
SNP: single-nucleotide polymorphism;
TDT: transmission disequilibrium test;
TGF: transforming growth factor;
TF: transcription factor;

## Acknowledgement

We would like to thank Marc Symons, Patricia Mongini, Kim Simpfendorfer, Jan Freudenberg and Ping Ao for discussion and suggestions. SM thanks Dr. Serena McCalla for support and guidance throughout the project, and thanks the financial support from the FIMR Summer Intern Program. WT and AS thank the support from the Robert S Boas Center for Genomics and Human Genetics.

## Appendix More information on the 17 craniosynostosis genes

The following table summarizes these information on the extra 17 craniosynostosis genes: (1) gene name and the corresponding OMIM number; (2) chromosome band location (the genes are ordered according to this column); (3) information about mutation, such as whether it’s de novo, whether it’s non-synonymous (missense) single-nucleotide polymorphism (nsSNP), or copy number variation (CNV), whether it’s autosomal dominant (AD) or autosomal recessive (AR) (question mark indicates the information is obtained from OMIM). There are two complex inheritance detection methods using many patients: one is TDT (transmission disequilibrium test) applied to many family trios (for BBS9), another is burden test of rare variance (SPRY1, SPRY4, SMURF1) – these methods are very different from the Mendelian gene detection methods. (4) information about phenotype, such as syndrome name (with OMIM number) for syndromic CRS, and suture location for non-syndromic CRS; (5) number of patients with the mutation described. For the two complex inheritance detection methods, the number of patients is indicated by a non-specific *n*; (6) the publication in which the gene mutation is studied.

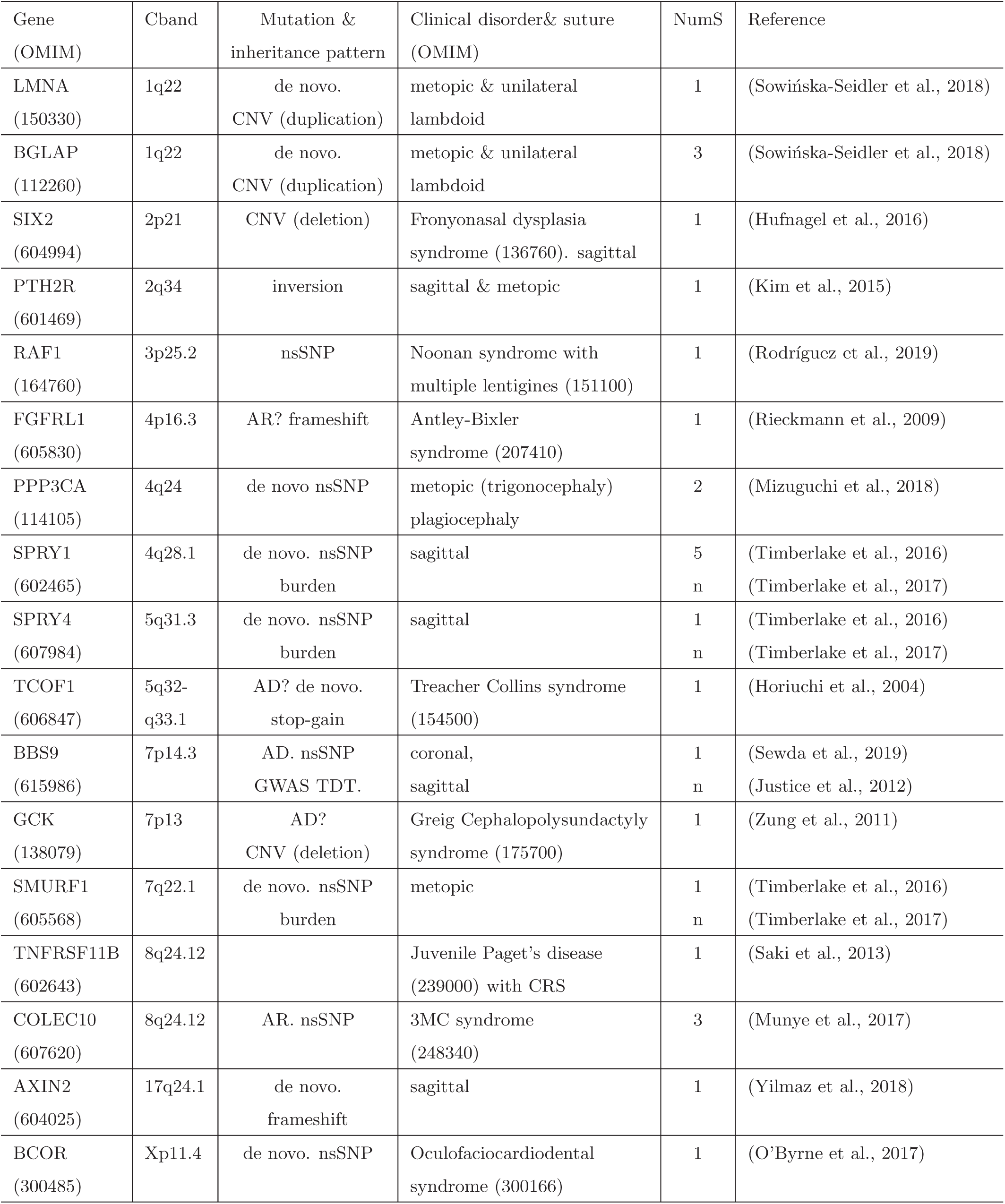

## Notes

#### Summary of Updates

A person's name in Acknowledgment was spelled incorrectly.

